# How does external lateral stabilization constrain normal gait, apart from improving medio-lateral gait stability?

**DOI:** 10.1101/2020.04.14.040535

**Authors:** Mohammadreza Mahaki, Trienke IJmker, Han Houdijk, Sjoerd Matthijs Bruijn

## Abstract

**Background:** The effect of external lateral stabilization on medio-lateral gait stability has been investigated previously. However, existing lateral stabilization devices not only constrains lateral motions, but also transverse and frontal pelvis rotations. This study aimed to investigate the effect of external lateral stabilization with and without constrained transverse pelvis rotation on mechanical and metabolic gait features.

**Methods:** We undertook 2 experiments with eleven and ten young adult subjects, respectively. Experiment 2 supplemented experiment 1, as it considered several potential confounding factors in the design and set-up of experiment 1. Kinematic, kinetic, and breath-by-breath oxygen consumption data were recorded during 3 walking conditions (normal walking (Normal), lateral stabilization with (Free) and without transverse pelvis rotation (Restricted)) and at 3 speeds (0.83, 1.25, and 1.66 m/s) for each condition.

**Results:** External lateral stabilization significantly reduced the amplitudes of the transverse and frontal pelvis rotations, medio-lateral pelvis displacement, transverse thorax rotation, arm swing, and step width. The amplitudes of free vertical moment, anterior-posterior and vertical pelvis displacements, step length, and energy cost were not significantly influenced by external lateral stabilization. The removal of transverse pelvis rotation restriction by our experimental set-up resulted in significantly higher transverse pelvis rotation, although it remained significantly less than Normal condition. In concert, concomitant gait features such as transverse thorax rotation and arm swing were not significantly influenced by our new set-up.

**Conclusion:** Existing lateral stabilization set-ups not only constrain medio-lateral motions (i.e. medio-lateral pelvis displacement), but also constrains other movements such as transverse and frontal pelvis rotations, which leads to several other gait changes such as reduced transverse thorax rotation, and arm swing. Our new setup allowed for more transverse pelvis rotation, however, this did not result in more normal pelvis rotation, arm swing, etc. Hence, to provide medio-lateral support without constraining other gait variables, more elaborate set-ups are needed. Unless such a set-up is realized the observed side effects need to be taken into account when interpreting the effects of lateral stabilization as reported in previous studies.

## 1. Introduction

Gait stability means keeping the body center of mass within the boundaries of the continuously changing base of support in the face of perturbations [1, 2]. Interactions of passive dynamics and active control are thought to be used by central nervous system to control gait stability [1, 2]. Different stabilizing strategies such as ankle [3], foot placement [2, 4-6], hip [7, 8], and push-off [8] strategies are used to control gait stability. To better understand the control of gait stability, external lateral stabilization devices have been used to manipulate medio-lateral stability [9-13, 4, 14]. It has been reported that external lateral stabilization reduces medio-lateral center of mass displacement [10] leading to lower need to control medio-lateral stability through the medio-lateral foot placement [10, 11, 4]. Some studies also reported that external lateral stabilization reduces metabolic energy cost of walking [10, 11, 14].

Existing Lateral stabilization devices not only constrain medio-lateral motions, but also transverse and frontal pelvis rotations as well as vertical pelvis displacement [11, 12]. During normal walking, the leg has to produce forces and moments to compensate for rotations along the longitudinal axis, and the arms contribute to this by having angular momentum in the opposite direction. Constraints of lateral external stabilization on transverse pelvis rotation may take away the need for the leg to produce such forces, and for the arms to counteract angular momentum produced by the swing leg [15]. In line with this, it has been reported that constrained pelvis rotation reduces the energy transfer from lower to upper extremities and subsequently reduces arm swing [16]. Transverse pelvis rotation also reduces vertical center of mass displacement [17] and contributes to step length, but only minimally [18]. Thus, apart from providing medio-lateral gait stability, lateral stabilization devices with constrained transverse pelvis rotation might limit the need to control angular momentum, reduces arm swing, and less likely changes the vertical center of mass displacement and step length. These potentially unwanted effects might change the interpretation of results reported by previous studies [10, 11, 14]. For example, the reduced energy cost in stabilized walking could be due to the aforementioned gait pattern constrains, rather than a reduced need to control medio-lateral gait stability [10, 11, 14].

The effect of constrained transverse pelvis rotation could be elucidated by adapting the standard lateral stabilization set-up to allow free pelvis rotations. Thus, we used two lateral stabilization conditions (with and without transverse pelvis rotation restriction) to determine how lateral stabilization set-ups constrain transverse pelvic rotation and concomitant gait features. We explored the direct effects of lateral stabilization on the amplitudes of the transverse and frontal pelvis rotations as well as the amplitudes of anterior-posterior and vertical pelvis displacements. Secondarily, we explored the indirect effects of the lateral stabilization on the amplitudes of transverse thorax rotation, arm swing, free vertical moment, and step length. To investigate the effects of lateral stabilization on medio-lateral gait stability, the effect of lateral stabilization on medio-lateral pelvis displacement (as a direct effect of lateral stabilization) as well as step width and energy cost (as two indirect effects of lateral stabilization) were explored^1^.

## 2. Methods

Experiments 1 and 2 were performed to test the effect of external lateral stabilization with and without constrained transverse pelvic rotation on the gait pattern (Figures 1. & 2.). Experiment 2 supplemented experiment 1, as it took into account several potential confounding factors in the experimental design or set-up. Moreover, additional experiments were performed to provide a detailed description of the characteristics of our set-up and the forces acting in it, which can be found in the Supplementary Material.

**Fig 1.**
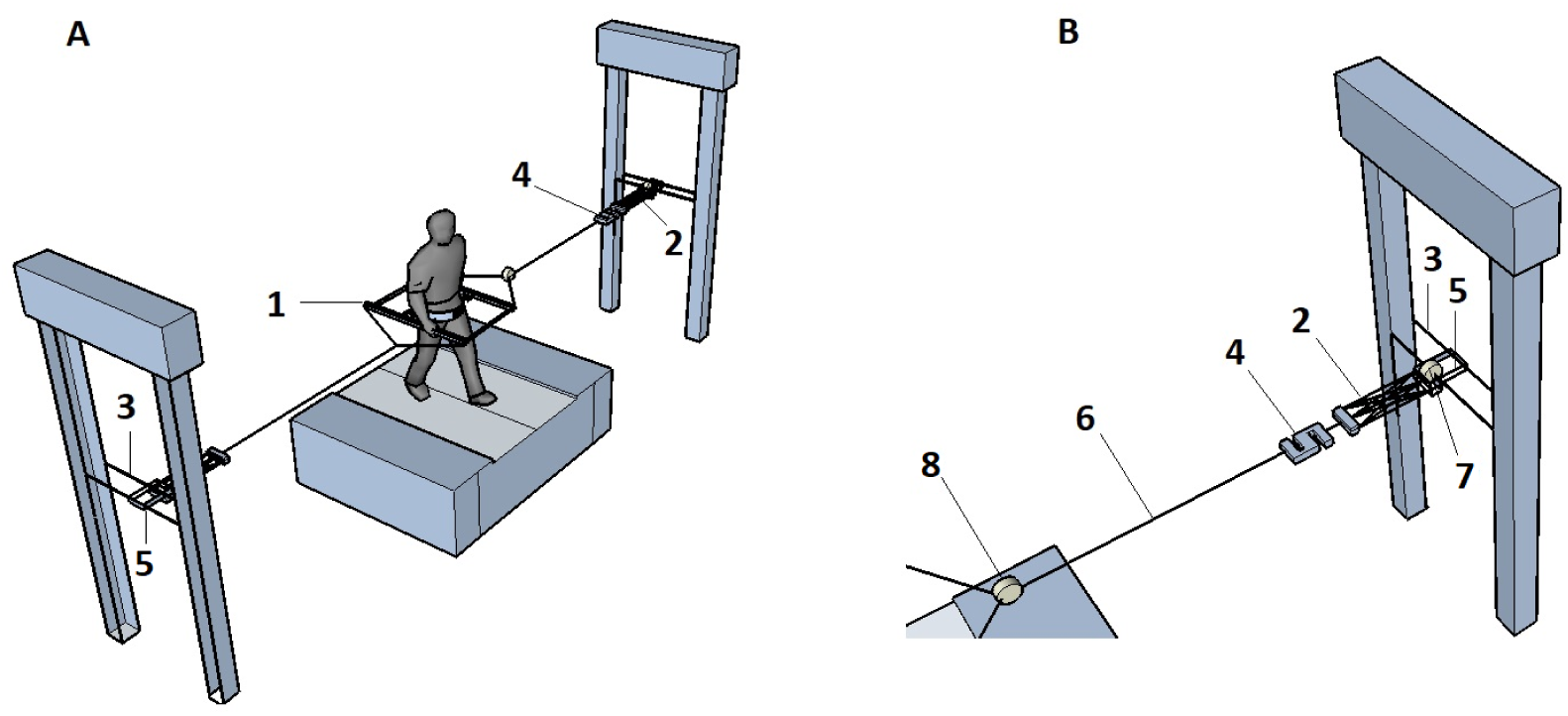
Schematic representation of experimental set up used in experiments 1 and 2. Inset (B) shows the stabilization in more detail. (1) Frame; (2) springs; (3) height-adjustable horizontal cart; (4) transducer (only used in additional experiments as reported in the supplementary material); (5) ball-bearing trolley freely moving in AP direction; (6) rope attached to frame; and (7 & 8) kinematics markers on the trolley and connection point between the springs and frame(only used in additional experiments as reported in the supplementary material).

**Fig 2.**
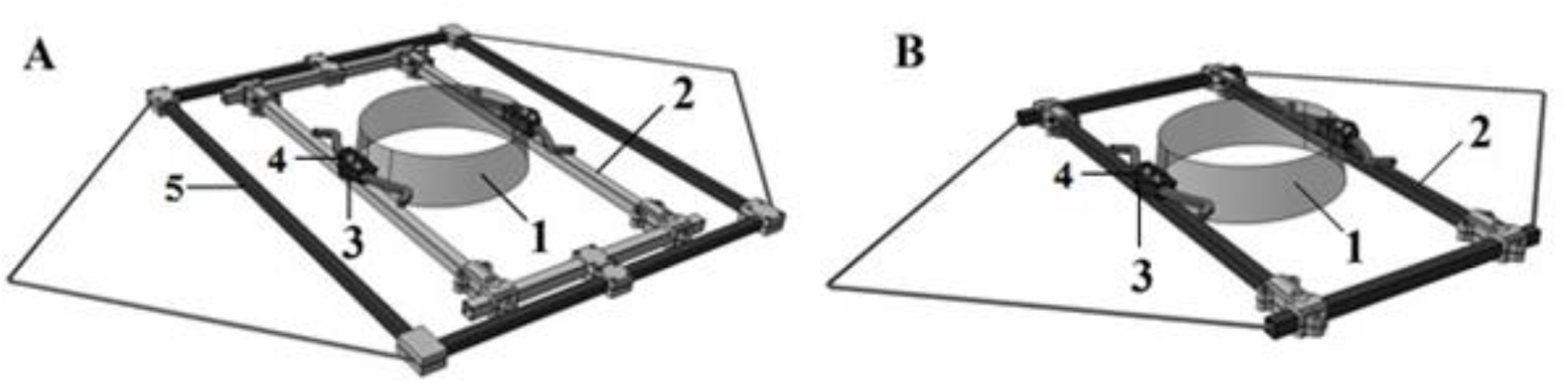
Schematic representation of the lateral stabilization frames used in experiments 1 (A) and 2 (B). (1) waist belt (we attached kinematic markers here); (2) inner frame which moves inside the outer frame to allow frontal pelvic rotation in experiment 1; (3) slider between waist belt and inner frame to allow transverse pelvic rotation; (4) Two screws resisting the sliders to work (5) outer frame allowing normal arm swing.

### 2.1 Experiment 1

#### 2.1.1 Participants

After signing the informed consent, a sample of 11 subjects (5 men and 6 women, age: 27.5 ± 2.4 years, mass: 62.6 ± 8.7 kg, and height: 1.72 ± 0.13 m (mean ± sd)) participated in this study, which was approved by the local ethics committee.

#### 2.1.2 Experimental set-up

A frame (weight = 4.5 kg) was used for the lateral stabilization. The frame was attached through a belt around the waist (Figure 2. A) and connected to bilateral springs of 1260 N m^-1^, which were connected to a rail placed at the height of the pelvis of the participant. This allowed free movements of the frame in the anterior-posterior direction. The frame included an inner and an outer frame, which were attached to each other and provided a free rotational degree of motion in the frontal plane between pelvis and frame. The distance between the inner and outer frames allowed normal arm swing during walking. In one condition, the pelvis was restricted from rotating in the transverse plane. In another condition, participants could rotate their pelvis with minimal friction between the waist belt and horizontal sliders on athe inner frame^1^.

#### 2.1.3 Experimental protocol

Participants were measured in three conditions (normal walking (Normal), lateral stabilization with transverse pelvis rotation restriction (Restricted), lateral stabilization without transverse pelvis rotation restriction (Free)). All conditions were executed at 3 speeds (0.83, 1.25, and 1.66 m/s). Each trial lasted 5 minutes, and trials were separated by a resting period of approximately 2 minutes. Conditions were randomized following a 2-step randomization procedure: For each participant, first the conditions were randomized and then speeds were randomized. No familiarization was provided and participants were instructed to walk normally during stabilized walking.

### 2.2 Experiment 2

#### 2.2.1 Participants

For experiment 2, a sample of 10 subjects (6 men and 4 women, age: 28.2 ± 3.7 years, mass: 70.3 ± 6.7 kg, and height: 1.77 ± 0.07 m (mean ± sd)) participated. Two of these participants also took part in experiment 1. Experiment 2 was executed under the same ethical approval as experiment 1.

#### 2.2.2 Experimental set-up

The frame in experiment 2 had no outer frame (Figure 2. B), thus had a reduced weight (weight =1.5 kg) compared to the frame used in experiment 1. Two stiff ropes attached to the frame on either side, joined each other at 0.5 m from the frame, providing space for free arm swing. From this junction, springs were attached to the springs of 1260 N/m and subsequently to the rail which allowed free movements in the anterior-posterior direction. This frame did not have a free degree of motion in frontal plane between pelvis and frame, thus restricted frontal pelvis rotation.

#### 2.2.3 Experimental protocol

The protocol for experiment 2 was equal to experiment 1, except that participants were familiarized with walking in each condition for about 2 minutes. Data collection started 10 minutes after the end of the familiarization protocol. Participants were instructed to walk normally during stabilized walking.

### 2.4 Instrumentation

During experiments 1 and 2, kinematic data was obtained from an Optotrak motion analysis system (Northern Digital Inc, Ontario, Canada), sampled at 100 samples/s. Clusters of three infrared markers were attached to the thorax, the pelvis, the waist belt of the frame, the left and right arms and the heels. We also obtained kinetic data from the force plates embedded in the treadmill (ForceLink b.v., Culemborg, the Netherlands), sampled at 200 samples/s in experiment 2. During all experiments, participants wore a mask and breath-by-breath oxygen consumption was obtained using a pulmonary gas exchange system (Cosmed K4b^2^, Cosmed, Italy).

### 2.5 Data processing

All our data and codes used to process the data for both experiments can be found at https://surfdrive.surf.nl/files/index.php/s/zZzjprWosewDaYf.

The direct effects of lateral stabilization were explored by calculating the amplitudes of transverse and frontal pelvis rotations as well as the amplitudes of medio-lateral, anterior-posterior and vertical pelvis displacements. To explore the indirect effects of lateral stabilization, the amplitudes of transverse thorax rotation, left and right arm swings, free vertical moment, left and right step lengths, step width and energy cost were calculated.

Kinematic data from the Optotrak system were not filtered. The displacement of three markers attached to each segment were averaged to quantify segment displacement. Kinematic data of heel markers was used to define the gait cycles. Each gait cycle was defined from left heel strike to subsequent left heel strike. Orientation of each segment was determined by a local anatomical reference frame constructed using three segment markers [19]. Using Euler angles, the time series of transverse and frontal pelvis rotations, transverse thorax rotation and arm swing were calculated from the segment orientation matrixes. The time series of angular (i.e. transverse and frontal pelvis rotations, transverse thorax rotation, and arm swing) and displacement (i.e. medio-lateral and vertical pelvis displacements) variables were time normalized to 0-100% for each gait cycle. The amplitudes of these variables were calculated as the differences between maximum and minimum angles/displacements per gait cycle and then median of amplitudes over gait cycles for each trial. The amplitude of the anterior-posterior pelvis displacement was calculated as the differences between maximum and minimum displacements over a walking trial. For arm swing, we calculated the average over arms since nonsignificant differences were found between left and right arm.

Ground reaction force data, collected in experiment 2, were filtered with a 5 Hz cut-off frequency (2nd order, bidirectional Butterworth digital filter). The time series of ground reaction forces were time normalized to 0-100% for each gait cycle. The vertical ground reaction moment (also referred to as ‘free vertical moment’, e.g. Li et al. 2001[20]) was calculated as the moment around a vertical axis caused by the interaction between the feet and the floor. The amplitude of free vertical moment was calculated as the differences between maximum and minimum vertical ground reaction moments per gait cycle and then the median of amplitudes over gait cycle for each trial. Step length and step width were defined as the median of the distances between anterior-posterior and medio-lateral foot placements at heel strike, respectively. For the step length, we calculated the average over legs since nonsignificant differences were found between left and right step lengths.

To evaluate the effect of lateral stabilization on energy cost, net energy cost was calculated. Oxygen uptake (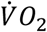; ml/min), carbon dioxide production (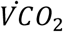;ml/min), and respiratory exchange ratio (RER) were determined with the pulmonary gas exchange system. The metabolic rate reached a plateau within the 5-minutes trial, as was confirmed through visual inspection. To be consistent with literature and to ensure that our outcome was not influenced by any potential noise, respiratory gases were averaged over the last three minutes of each trial. We calculated Gross metabolic rate (E_gross_; J/min) using the following equation [21]:

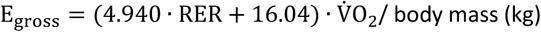

Resting metabolic rate, determined during seated position for 5 min prior to the trials, was subtracted from gross metabolic rate to calculate net metabolic rate during walking. To calculate net energy cost (J/kg/m), net metabolic rate was divided by walking speed (m/min).

### 2.6 Statistical analysis

Repeated measures ANOVA with Condition ([Normal, Free, and Restricted]) and Speed ([0.83, 1.25, and 1.66 m/s]) as factors was used to test for the effects of external lateral stabilization with and without transverse pelvic rotation and walking speed on all direct and indirect outcomes. After finding a significant main effect of Condition, a post-hoc test with Bonferroni correction was applied to determine the differences between conditions. We assessed the Condition × Speed interaction, to test for the differences in the effects of external lateral stabilization with and without transverse pelvis rotation on aforementioned parameters between different walking speeds. For F-test, 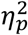 was reported as effect size. Level of significance for all statistical analyses was set at p < 0.05.

## 3. Results

### 3.1 Experiment 1

Direct effects of lateral stabilization were found in the amplitudes of transverse pelvis rotation (Figure 3. A), frontal pelvis rotation (Figure 3. C), and medio-lateral pelvis displacement (Figure 4. A), which were significantly influenced by Condition (Table 1.). Post-hoc analyses showed that in Free and Restricted conditions, the amplitudes of transverse pelvis rotation, frontal pelvis rotation, and medio-lateral pelvis displacement were significantly reduced when compared to the Normal condition (Table 1.). However, the differences of influenced gait variables between Free and Restricted conditions were not significant (Table 1.). The amplitudes of anterior-posterior and vertical pelvis displacements (Figure 4. C & E) were not significantly influenced by Condition (Table 1.).

**Table 1.**
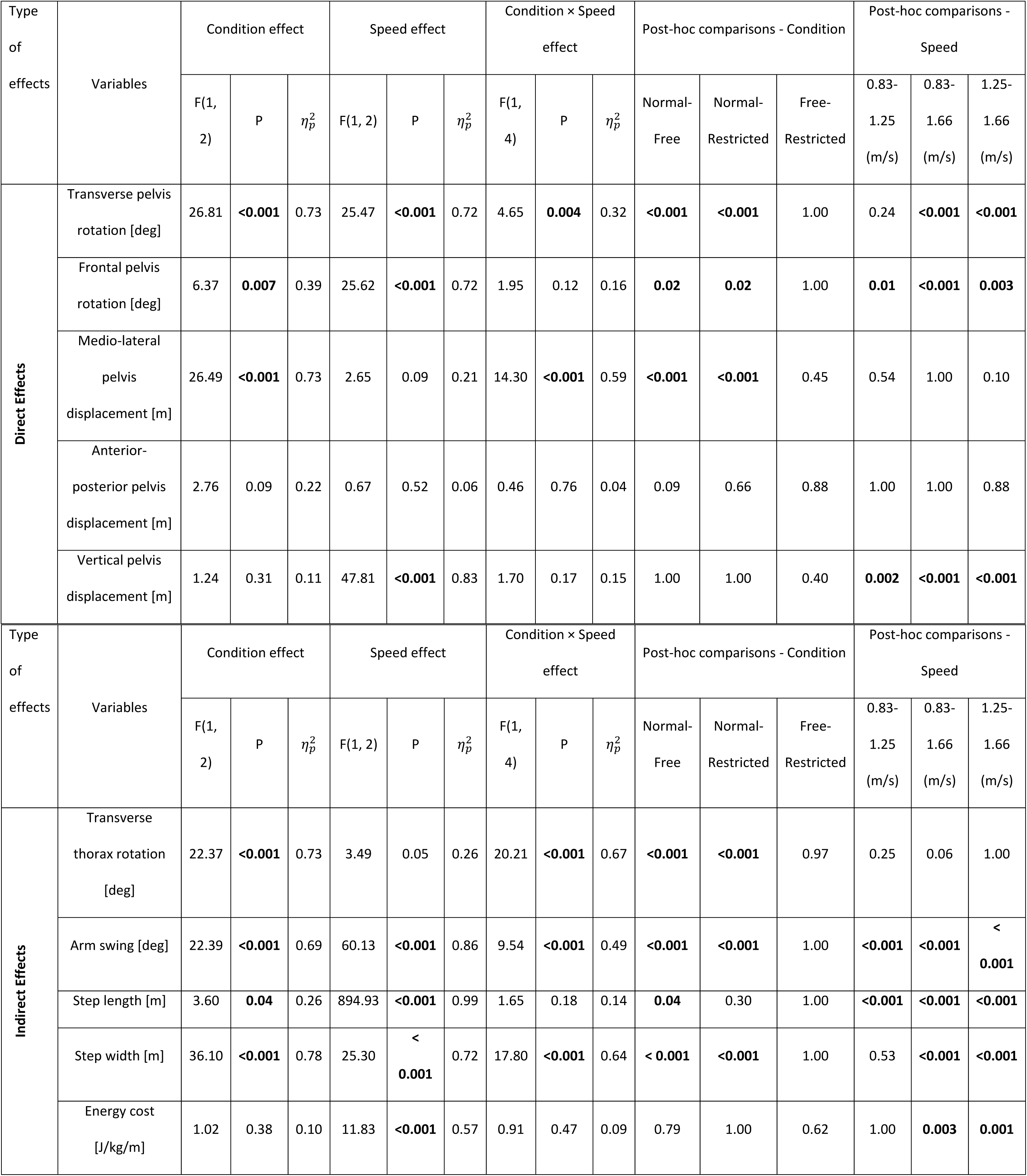
The direct and indirect effect of external lateral stabilization on mechanical and physiological gait variables in experiment 1.

| Type of effects | Variables | Condition effect |  |  | Speed effect |  |  | Condition × Speed effect |  |  | Post-hoc comparisons - Condition |  |  | Post-hoc comparisons - Speed |  |  |
| --- | --- | --- | --- | --- | --- | --- | --- | --- | --- | --- | --- | --- | --- | --- | --- | --- |
| | | F(1, 2) | P | $\eta_p^2$ | F(1, 2) | P | $\eta_p^2$ | F(1, 4) | P | $\eta_p^2$ | Normal-Free | Normal-Restricted | Free-Restricted | 0.83-1.25 (m/s) | 0.83-1.66 (m/s) | 1.25-1.66 (m/s) |
| Direct Effects | Transverse pelvis rotation [deg] | 26.81 | **<0.001** | 0.73 | 25.47 | **<0.001** | 0.72 | 4.65 | **0.004** | 0.32 | **<0.001** | **<0.001** | 1.00 | 0.24 | **<0.001** | **<0.001** |
|  | Frontal pelvis rotation [deg] | 6.37 | **0.007** | 0.39 | 25.62 | **<0.001** | 0.72 | 1.95 | 0.12 | 0.16 | **0.02** | **0.02** | 1.00 | **0.01** | **<0.001** | **0.003** |
|  | Medio-lateral pelvis displacement [m] | 26.49 | **<0.001** | 0.73 | 2.65 | 0.09 | 0.21 | 14.30 | **<0.001** | 0.59 | **<0.001** | **<0.001** | 0.45 | 0.54 | 1.00 | 0.10 |
|  | Anterior-posterior pelvis displacement [m] | 2.76 | 0.09 | 0.22 | 0.67 | 0.52 | 0.06 | 0.46 | 0.76 | 0.04 | 0.09 | 0.66 | 0.88 | 1.00 | 1.00 | 0.88 |
|  | Vertical pelvis displacement [m] | 1.24 | 0.31 | 0.11 | 47.81 | **<0.001** | 0.83 | 1.70 | 0.17 | 0.15 | 1.00 | 1.00 | 0.40 | **0.002** | **<0.001** | **<0.001** |
| | | F(1, 2) | P | $\eta_p^2$ | F(1, 2) | P | $\eta_p^2$ | F(1, 4) | P | $\eta_p^2$ | Normal-Free | Normal-Restricted | Free-Restricted | 0.83-1.25 (m/s) | 0.83-1.66 (m/s) | 1.25-1.66 (m/s) |
| Indirect Effects | Transverse thorax rotation [deg] | 22.37 | **<0.001** | 0.73 | 3.49 | 0.05 | 0.26 | 20.21 | **<0.001** | 0.67 | **<0.001** | **<0.001** | 0.97 | 0.25 | 0.06 | 1.00 |
|  | Arm swing [deg] | 22.39 | **<0.001** | 0.69 | 60.13 | **<0.001** | 0.86 | 9.54 | **<0.001** | 0.49 | **<0.001** | **<0.001** | 1.00 | **<0.001** | **<0.001** | **< 0.001** |
|  | Step length [m] | 3.60 | **0.04** | 0.26 | 894.93 | **<0.001** | 0.99 | 1.65 | 0.18 | 0.14 | **0.04** | 0.30 | 1.00 | **<0.001** | **<0.001** | **<0.001** |
|  | Step width [m] | 36.10 | **<0.001** | 0.78 | 25.30 | **< 0.001** | 0.72 | 17.80 | **<0.001** | 0.64 | **< 0.001** | **<0.001** | 1.00 | 0.53 | **<0.001** | **<0.001** |
|  | Energy cost [J/kg/m] | 1.02 | 0.38 | 0.10 | 11.83 | **<0.001** | 0.57 | 0.91 | 0.47 | 0.09 | 0.79 | 1.00 | 0.62 | 1.00 | **0.003** | **0.001** |

**Fig 3.**
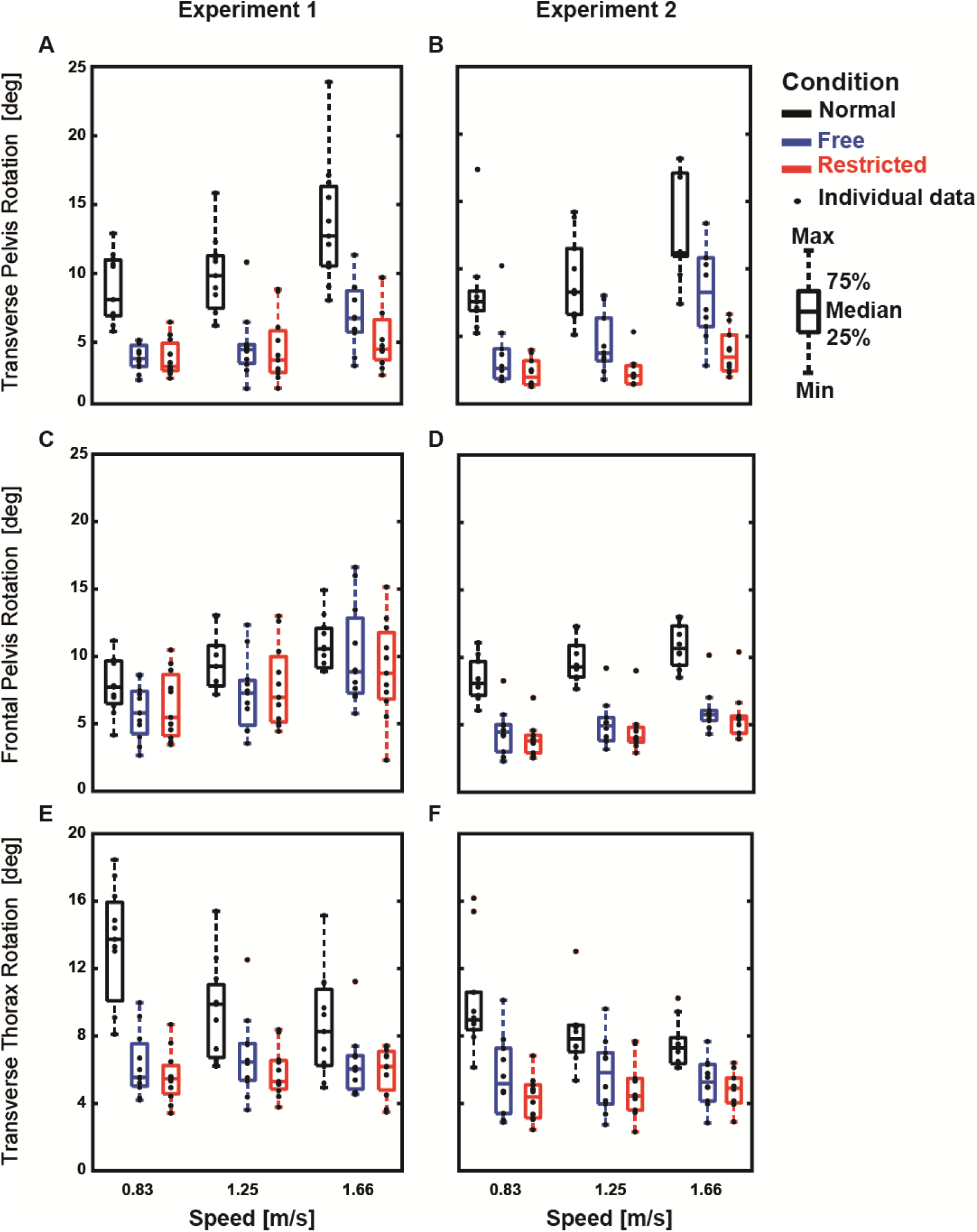
Group average limb kinematics (median ± 25^th^ percentile) at each walking speed in the Normal (black), stabilized conditions without transverse pelvis rotation restriction (blue) and with transverse pelvis rotation restriction (red) for transverse pelvis rotation **(A & B)** and frontal pelvis rotation **(C & D)** as well as transverse thorax rotation **(E & F)** in experiments 1 and 2.

**Fig 4.**
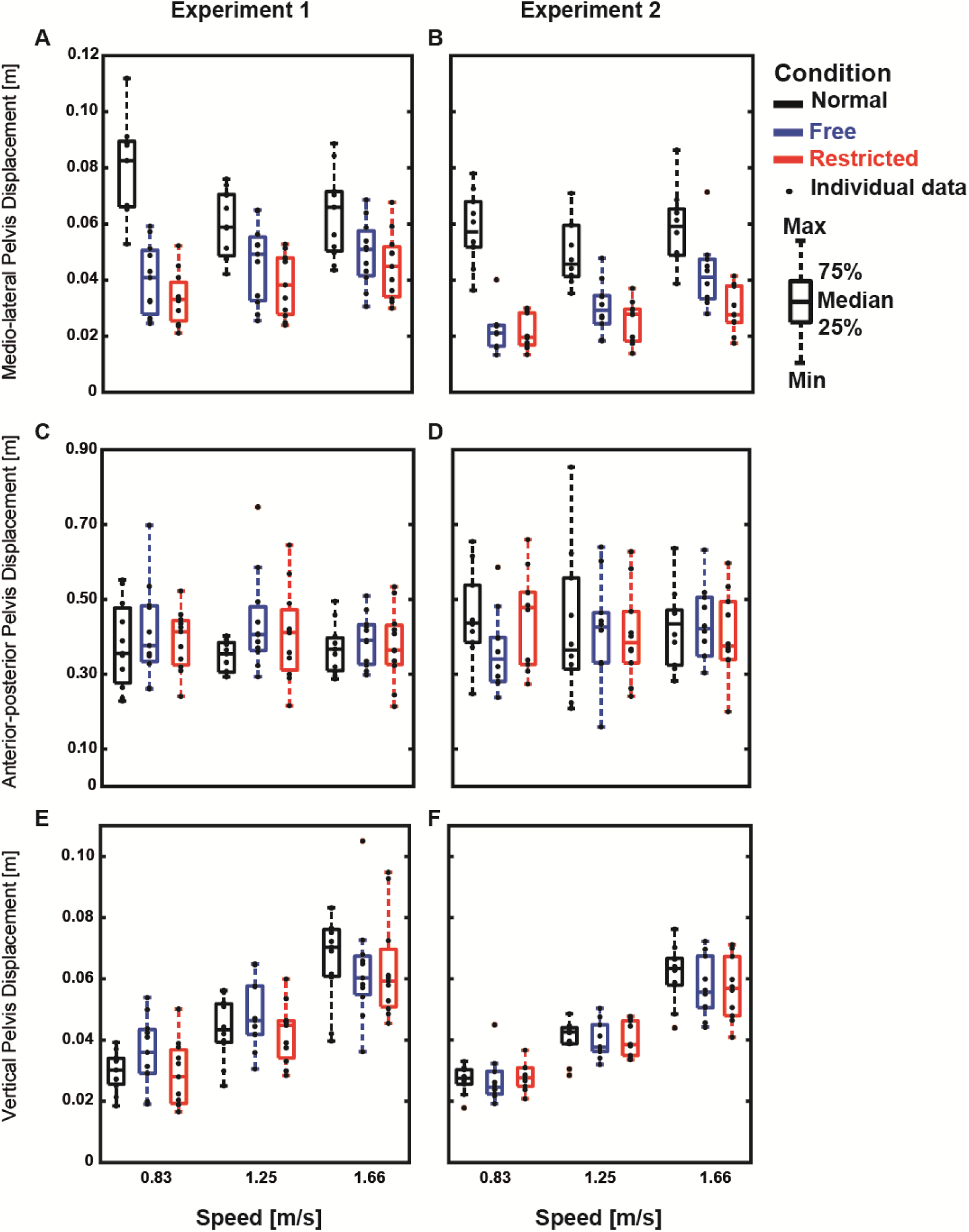
Group average limb kinematics (median ± 25^th^ percentile) at each walking speed in the Normal (black), stabilized conditions without transverse pelvis rotation restriction (blue) and with transverse pelvis rotation restriction (red) for medio-lateral **(A & B)**, anterior-posterior **(C & D)**, and vertical pelvis displacements **(E & F)** in experiments 1 and 2.

Indirect effects of lateral stabilization were found in the amplitudes of transverse thorax rotation (Figure 3. E), arm swing (Figure 5. A), step length (Figure 6. A) and step width (Figure 6. C), which were significantly influenced by Condition. Energy cost (Figure 7. A) was not significantly influenced by Condition (Table 1.). Post-hoc analyses showed that in Free and Restricted conditions, the influenced gait variables significantly reduced when compared to the Normal condition (Table 1.). However, the differences of influenced gait variables between Free and Restricted conditions were not significant (Table 1.).

**Fig 5.**
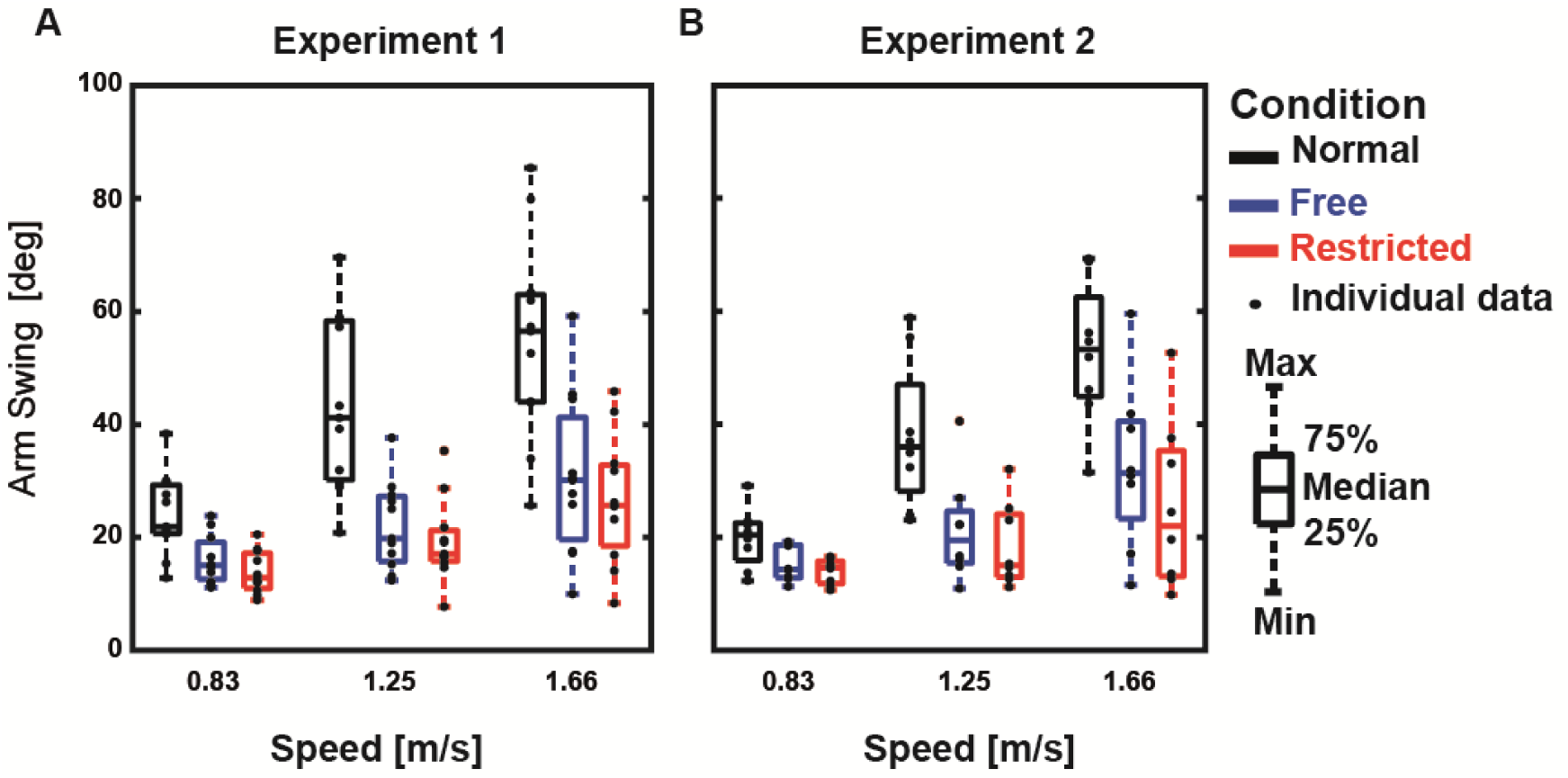
Group average limb kinematics (median ± 25^th^ percentile) at each walking speed in the Normal (black), stabilized conditions without transverse pelvis rotation restriction (blue) and with transverse pelvis rotation restriction (red) for arm swing in experiments 1 **(A)** and 2 **(B)**.

**Fig 6.**
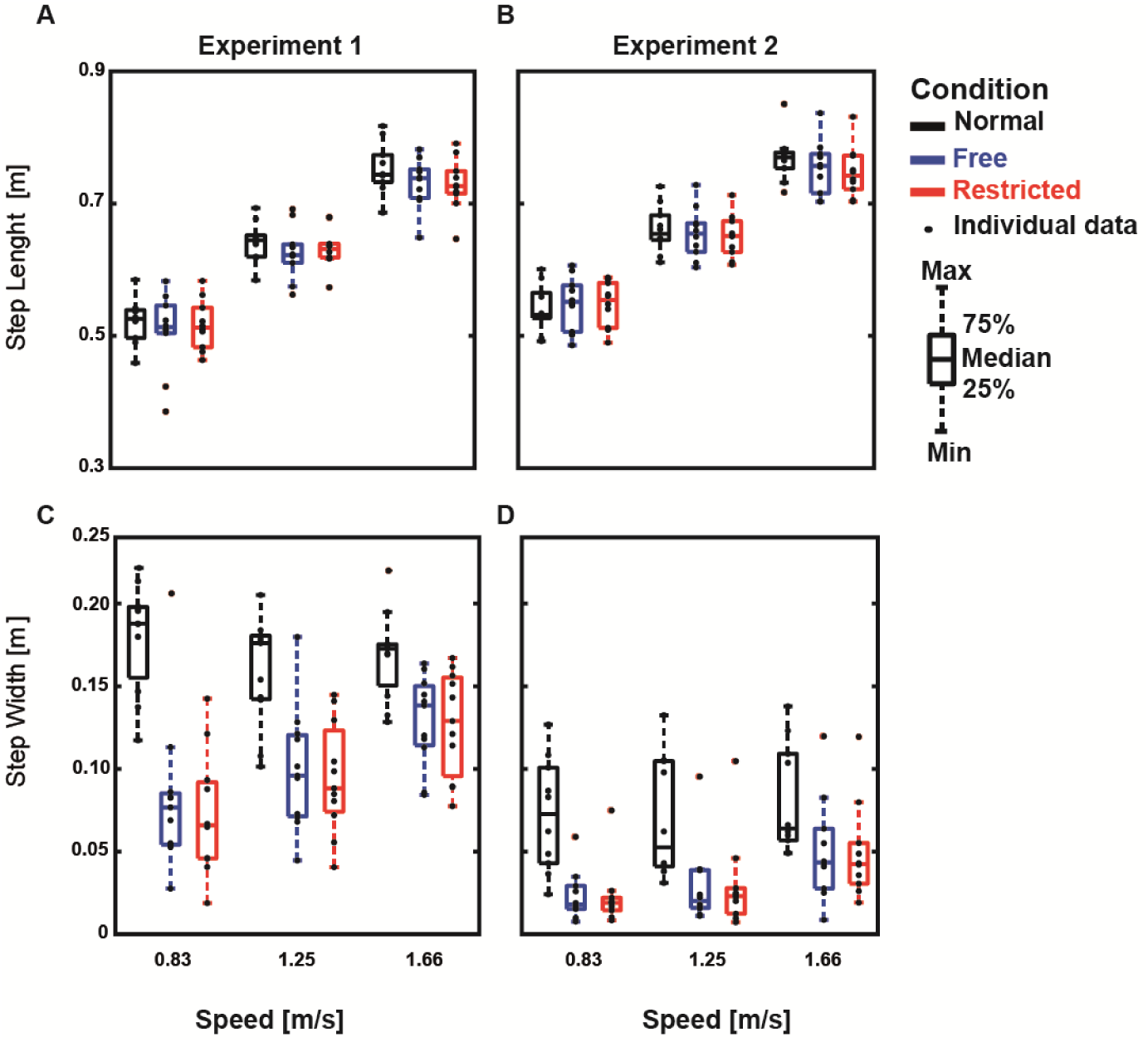
Group average limb kinematics (median ± 25^th^ percentile) at each walking speed in the Normal (black), stabilized conditions without transverse pelvis rotation restriction (blue) and with transverse pelvis rotation restriction (red) for step length **(A & B)** and step width **(C & D)** in experiments 1 and 2.

**Fig 7.**
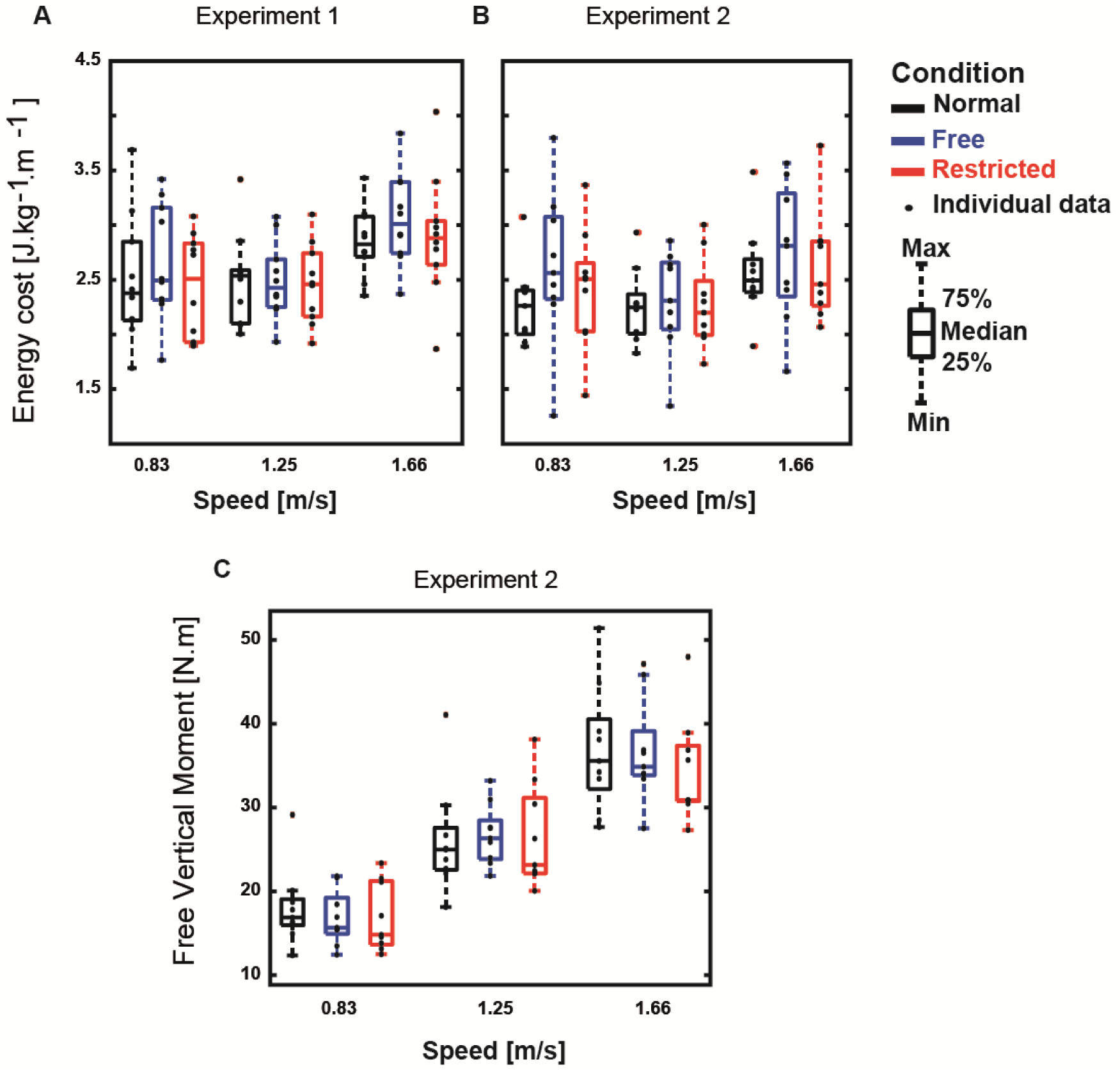
Group average (median ± 25^th^ percentile) at each walking speed in the Normal (black), stabilized conditions without transverse pelvis rotation restriction (blue) and with transverse pelvis rotation restriction (red) for energy cost of walking in experiments 1 **(A)** and 2 **(B)**, as well as for free vertical moment in experiment 2 **(C).**

At faster walking speeds, the amplitudes of transverse and frontal pelvis rotations (Figure 3. A & C), arm swing (Figure 5. A), vertical pelvis displacement (Figure 4. E), step length (Figure 6. A), step width (Figure 6. C), and energy cost (Figure 7. A) increased (Speed effect and related post-hoc analyses; Table 1.). The increases of transverse pelvis rotation (Figure 3. A) and arm swing (Figure 5. A) by increasing walking speed were shown to be more prominent in Normal condition when compared to the Free and Restricted conditions (Condition × Speed effects; Table 1.). No significant differences of transverse pelvis rotation and arm swing by increasing walking speed were found between Free and Restricted conditions (Condition × Speed effects; Table 1.).

### 3.2 Experiment 2

Direct effects of lateral stabilization were found in the amplitudes of transverse pelvis rotation (Figure 3. B), frontal pelvis rotation (Figure 3. D), and medio-lateral pelvis displacement (Figure 4. B), which were significantly influenced by Condition (Table 2.). Post-hoc analyses showed that the amplitude of transverse pelvis rotation was significantly higher in Normal than in Free and in Free than in Restricted condition (Figure 3. B and Table 2.). In Free and Restricted conditions, frontal pelvis rotation and medio-lateral pelvis displacement were significantly reduced when compared to the Normal condition (Table 2.). Like in experiment 1 the amplitudes of anterior-posterior and vertical pelvis displacements (Figure 4. D & F) were not significantly influenced by Condition (Table 2.).

**Table 2.** The direct and indirect effect of external lateral stabilization on mechanical and physiological gait variables in experiment 2.

| Type of effects | Variables | Condition effect |  |  | Speed effect |  |  | Condition × Speed |  |  | Post-hoc comparisons - Condition |  |  | Post-hoc comparisons - Speed |  |  |
| --- | --- | --- | --- | --- | --- | --- | --- | --- | --- | --- | --- | --- | --- | --- | --- | --- |
| | | F(1, 2) | P | $\eta_p^2$ | F(1, 2) | P | $\eta_p^2$ | F(1, 4) | P | $\eta_p^2$ | Normal - Free | Normal-Restricted | Free-Restricted | 0.83-1.25 (m/s) | 0.83-1.66 (m/s) | 1.25-1.66 (m/s) |
| Direct Effects | Transverse pelvis rotation [deg] | 33.07 | < **0.001** | 0.79 | 24.77 | < **0.001** | 0.73 | 5.17 | **0.002** | 0.37 | < **0.001** | < **0.001** | **0.03** | 0.91 | < **0.001** | < **0.001** |
|  | Frontal pelvis rotation [deg] | 57.88 | < **0.001** | 0.87 | 29.24 | < **0.001** | 0.77 | 2.30 | 0.08 | 0.20 | < **0.001** | < **0.001** | 0.96 | **0.008** | < **0.001** | **0.002** |
|  | Medio-lateral pelvis displacement [m] | 50.11 | < **0.001** | 0.85 | 21.67 | < **0.001** | 0.71 | 8.51 | < **0.001** | 0.49 | < **0.001** | < **0.001** | 0.18 | 1.00 | < **0.001** | < **0.001** |
|  | Anterior-posterior pelvis displacement [m] | 0.49 | 0.62 | 0.05 | 0.02 | 0.98 | 0.002 | 1.59 | 0.20 | 0.15 | 1.00 | 1.00 | 1.00 | 1.00 | 1.00 | 1.00 |
|  | Vertical pelvis displacement [m] | 1.08 | 0.36 | 0.11 | 184.49 | < **0.001** | 0.95 | 4.39 | **0.005** | 0.33 | 0.55 | 0.85 | 1.00 | < **0.001** | < **0.001** | < **0.001** |
| | | F(1, 2) | P | $\eta_p^2$ | F(1, 2) | P | $\eta_p^2$ | F(1, 4) | P | $\eta_p^2$ | Normal - Free | Normal-Restricted | Free-Restricted | 0.83-1.25 (m/s) | 0.83-1.66 (m/s) | 1.25-1.66 (m/s) |
| Indirect Effects | Transverse thorax rotation [deg] | 28.77 | < **0.001** | 0.76 | 3.10 | 0.07 | 0.26 | 5.66 | **0.001** | 0.39 | < **0.001** | < **0.001** | 0.29 | 0.72 | 0.07 | 0.66 |
|  | Arm swing [deg] | 34.1 | < **0.001** | 0.83 | 25.6 | < **0.001** | 0.79 | 11.40 | **0.04** | 0.62 | < **0.001** | < **0.001** | 0.24 | **0.01** | < **0.001** | **0.005** |
|  | Free vertical moment [N.m] | 0.48 | 0.63 | 0.06 | 192.71 | < **0.001** | 0.96 | 1.69 | 0.18 | 0.18 | 1.00 | 1.00 | 1.00 | < **0.001** | < **0.001** | < **0.001** |
|  | Step length [m] | 3.04 | 0.07 | 0.25 | 547.52 | < **0.001** | 0.98 | 4.86 | **0.003** | 0.35 | 0.45 | 0.06 | 0.44 | < **0.001** | < **0.001** | < **0.001** |
|  | Step width [m] | 21.54 | < **0.001** | 0.71 | 13.29 | < **0.001** | 0.60 | 3.48 | **0.02** | 0.28 | < **0.001** | < **0.001** | 1.00 | 1.00 | < **0.001** | **0.002** |
|  | Energy cost [J/kg/m] | 1.456 | 0.263 | 0.15 | 8.15 | **0.004** | 0.51 | 2.72 | **0.047** | 0.25 | 0.33 | 1.00 | 0.95 | 0.31 | 0.11 | **0.003** |

Indirect effects of lateral stabilization were found in the amplitudes of transverse thorax rotation (Figure 3. F), arm swing (Figure 5. B), and step width (Figure 6. D), which were significantly influenced by Condition (Table 2.). Step length (Figure 6. B), free vertical moment (Figure 7. C), and energy cost (Figure 7. B) were not significantly influenced by Condition (Table 2.). Post-hoc analyses showed that the differences of transverse thorax rotation, arm swing, and step width between Free and Restricted conditions were not significant (Table 2.). In both Free and Restricted conditions, transverse thorax rotation, arm swing, and step width were significantly reduced when compared to the Normal condition (Table 2.).

At faster walking speeds, the amplitudes of transverse and frontal pelvis rotations (Figure 3. B & D), arm swing (Figure 5. B), medio-lateral and vertical pelvis displacements (Figure 4. B & F), step length (Figure 6. B), step width (Figure 6. D), free vertical moment (Figure 7. C) and energy cost (Figure 7. B) increased (Speed effect and related post-hoc analyses; Table 2.). The increase of transverse pelvis rotation with increasing of walking speed was more pronounced in the Normal condition than in the Free condition as well as in Free condition than in Restricted condition (Condition × Speed effect; Figure 3. B & Table 2.). The increases of arm swing, medio-lateral and vertical pelvis displacements, step length, and step width were more pronounced in the Normal condition when compared to the Free and Restricted conditions (Condition × Speed effect; Table 2.).

## 4. Discussion

We used external lateral stabilization with and without transverse pelvis rotation to investigate how existing lateral stabilization set-ups constrain gait apart from providing medio-lateral gait stability. Our results demonstrated that external lateral stabilization with and without transverse pelvis rotation not only constrains medio-lateral motions (i.e. the amplitudes of medio-lateral pelvis displacement and step width), but also constrains transverse and frontal pelvis rotations, which coincided with several other gait changes such as reduced transverse thorax rotation, and reduced arm swing which could be considered an indirect consequence of the restricted pelvis motions. While the removal of constraints in our experimental set-up led to increased transverse and frontal pelvis rotations, the amplitudes of anterior-posterior and vertical pelvis displacements, free vertical moment, and energy cost were not influenced by lateral stabilization. The increases of transverse pelvis rotation with increasing speed was shown to be more dominant in Free condition when compared to the Restricted condition. However, the increases of arm swing, medio-lateral and vertical pelvis displacements, step length, and step width with increasing speed were shown to be more dominant in Normal condition, but did not parallel the difference in pelvic rotation that was observed between both stabilized conditions.

Consistent with previous studies [10, 11, 13, 4], our results confirmed that external lateral stabilization provided medio-lateral gait stability, represented by a direct reduction of medio-lateral pelvis displacement which was accompanied by an indirect reduction of step width in stabilized condition. Consistent with Matsubara et al. [22], our results confirmed that external lateral stabilization did not constrain the amplitude of anterior-posterior pelvis displacement because bilateral springs were connected to the horizontal trolley which were capable to move freely and in-phase with the displacement of the center of mass in anterior-posterior direction. However, springs fixed in anterior-posterior direction used by previous studies [23, 10, 11, 14] may provide unwanted forces and assistance in the anterior-posterior direction (see Supplementary Material; Figure. S1) and subsequently decreases the net propulsive force generating by the subjects (cf. [24]). It has been reported that humans try to decrease levels of muscle activation during walking in force fields which has also been called “slacking” [25, 26]. Thus, in previous studies with fixed springs [23, 10, 11, 14], subjects may have benefited from slacking in the stabilized condition if they discovered that walking is less energetically costly at the back of the treadmill than at other spots. Here, unwanted anterior-posterior component forces pull the subject forward, potentially leading to lower levels of muscle activation and reduced propulsive force. Although some studies tried to minimize these potential unwanted effects by using long ropes to connect the springs to the subjects (i.e. 8.5 and 14.5 m [11, 14], respectively), other studies with shorter ropes (i.e. 3.0 and 4.0 m [23, 10], respectively) might have increased these effects and subsequently some of the energy cost savings of external lateral stabilization found in these studies may have been due to “slacking”.

Using two transverse sliders between waist belt and inner frame, our results showed that transverse pelvis rotation was less restricted by lateral stabilization in the Free condition than in the Restricted condition (i.e. the set-up used by previous studies) in experiment 2. The higher transverse pelvis rotation in the Free condition than in the Restricted condition was more pronounced with increasing walking speed. Our results also showed that frontal plane pelvis rotation was less restricted by lateral stabilization with the frame used in experiment 1 than with the frame used in experiment 2 (see Figure 3. C & D). The frame used in experiment 1 included an inner and an outer frames, which were attached to each other and provided a free rotational degree of motion in the frontal plane between pelvis and frame, however the frame used in experiment 2 did not have an outer frame and it was more similar to the frame used by previous studies [12, 13]. Therefore, it seems that our new set-ups restricted transverse and frontal pelvis rotations less than previous set-ups.

Although subjects were allowed to perform arm swing and they were instructed to walk as naturally as possible, our results showed reduced arm swing in stabilized conditions. It has been reported that the restriction of transverse pelvis rotation reduces the energy transfer between the lower and upper limb segments and subsequently deceases the amplitude of arm swing [16]. The reduced arm swing might be due to restricted transverse pelvis rotation in stabilized conditions since our new set-up did not lead to fully normal transverse pelvis rotation. However, arm swing was less restricted in the present study than those studies in which subjects were not allowed to swing their arms [10, 11] and they had to walk with their arms crossed (to avoid contacting the external stabilizer). Our results also showed that free vertical moment was not influenced by lateral stabilization. This indicates that the control of angular momentum was not limited in stabilized conditions [15]. Previous studies which provided lateral stabilization with completely constrained pelvis rotation and arm swing [10, 27] might have limited the need to control of angular momentum as hypothesized by [15]. It can be concluded that our set-up and the set-ups used by previous studies not only constrain medio-lateral motions (i.e. medio-lateral pelvis displacement and step width), but also transverse and frontal pelvis rotation, thereby indirectly leading to decreases in transverse thorax rotation and arm swing.

In contrast to some of previous studies [11, 14], we did not find a decrease in energy cost during stabilized walking. This appears to contradict the notion that medio-lateral stabilization is an active process which entails energy cost. However, there may be several explanations for this. Firstly, springs that are fixed in anterior-posterior direction as used by previous studies [11, 14] might provide unwanted forces and assistance in the anterior-posterior direction and might have increased some of the energy cost savings of external lateral stabilization. Secondly, it has been reported that with arm swing the effect of lateral stabilization on energy cost was slightly lower (a significant 3-4% reduction), compared to walking without arm swing (a significant 6-7% reduction) [14]. Therefore, arm swing restriction induced by set-up used by previous studies [10, 11] might also have increased some of the energy cost savings of lateral stabilization. Thirdly, providing more than 30 minutes habituation time, Ortega et al. [14] reported a significant 3-4% reduction of energy cost in stabilized condition. In contrast to Ortega et al. [14], a habituation time between 3-10 minutes was used in the current study and in the previous studies [10, 13] with the reported nonsignificant effect of lateral stabilization on energy cost. Therefore, not long enough habituation time to allow for full familiarization might be responsible for the inability of our set-up to reduce energy cost. Lastly, although the potential effects of the frame weight and it’s rotational inertia on energy cost were minimized by removing outer frame in the experiment 2, the frame used by IJmker et al. [12, 13] and our two experiments may have added additional energy cost and subsequently these factors might offset some of the reduced energy cost induced by external lateral stabilization. To draw a conclusion about these observed effects of lateral stabilization on energy cost, a meta-analysis on the available data could provide valuable insights.

The restriction of transverse and frontal pelvis rotations induced by the existing lateral stabilization set-ups leads to changes in the gait pattern beyond those that are targeted by this manipulation. Therefore, when interpreting the results of lateral stabilization studies, this limitation should be considered. To more accurately evaluate the effect of lateral stabilization on the control and energy cost of medio-lateral gait stability, future lateral stabilization studies are suggested to improve their set-ups and to refrain from restricting gait patterns such as transverse and frontal pelvis rotations, and anterior-posterior pelvis displacement.

## 5. Conclusion

External lateral stabilization set-ups do not only constrain the amplitudes of the medio-lateral motions, but also the amplitudes of the transverse and frontal pelvis rotations, leading to reduced transverse thorax rotation as well as arm swing. Although the removal of constraints by our experimental set-up led to increased transverse and frontal pelvis rotations, other concomitant gait variables such as transverse thorax rotation and arm swing still remained restricted in stabilized conditions. Therefore, future studies are recommended to take these limitations of lateral stabilization set-ups into consideration and to improve their set-ups.

## Acknowledgements

SMB was funded by a VIDI grant (016.Vidi.178.014) from the Dutch Organization for Scientific Research (NWO). MM was also founded by a grant from Ministry of Science, Research and Technology of Iran.

## Footnotes

1 Our initial research proposal can be found at https://osf.io/gkphs/.

1 The detailed description of frictions between waist belt and horizontal sliders as well as between ball-bearing trolley and cart can be found in the Supplementary Material.

## Notes

https://surfdrive.surf.nl/files/index.php/s/zZzjprWosewDaYf

https://osf.io/gkphs/

### Competing Interest Statement

The authors have declared no competing interest.

