## Supplementary Material file (Fig S1., Fig S2., and Fig S3.) for "How does external lateral stabilization constrain normal gait, apart from improving medio-lateral gait stability?"

### The estimation of anterior-posterior forces induced by bilateral springs

In the present study, bilateral springs were connected to the bilateral trolleys which were capable to move in the anterior-posterior direction (Free trolley; **Fig S1. A**). However, previous studies [1-4] have used fixed springs in the anterior-posterior direction (Fixed trolley; **Fig. S1. B**). To check if the movement of the trolley caused unwanted oscillations, which could cause unwanted anterior-posterior forces and assistance acting on the body, we performed an experiment with one subject. To check if the position of the subject on the treadmill caused unwanted anterior-posterior forces, the subject was asked to walk with and without extra movement in anterior-posterior direction. Two markers and a force transducer in series with the rope were used to collect kinematics and force data during walking at 1.25 (m/s) in all conditions. This experiment was performed with the frame which we used in experiment 2.

The results showed the displacements of the trolley and frame were in-phase during walking with Free trolley condition (**Fig S1. C**). In both conditions (i.e. Fixed and Free Trolley), the direction and magnitude of anterior-posterior forces were approximately similar, and quite small (< ± 5N) when subject was asked to walk without extra movement in anterior-posterior direction (Normal Walking; **Fig S1. E & F**). However, when the subject was asked to walk with extra movement in the anterior-posterior direction, the forces in anterior-posterior direction were smaller in the Free trolley condition (Extra anterior-posterior movements; **Fig S1. G & H**).

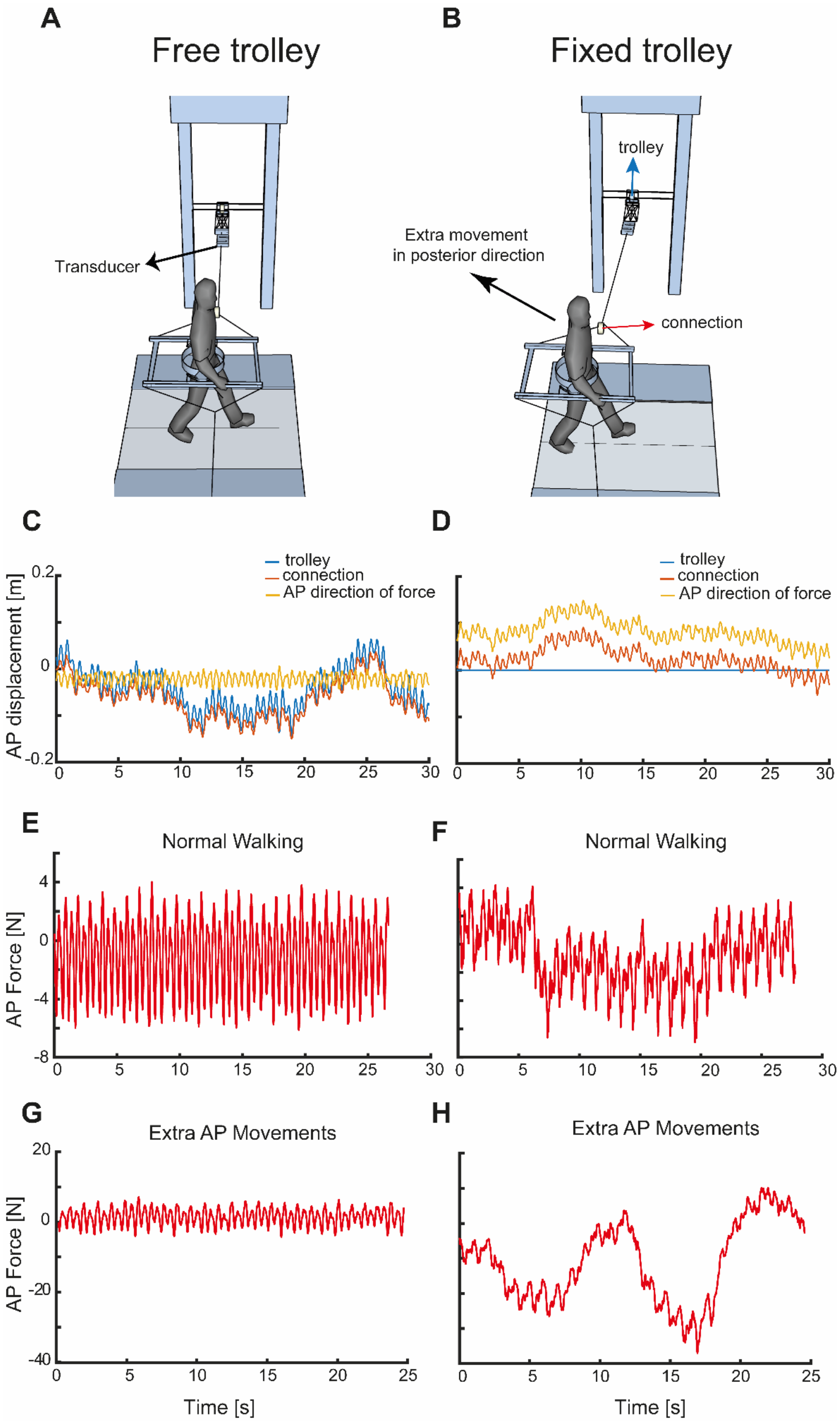


**Fig S1.** **(A)** Free trolley in the anterior-posterior direction represents the set-up used in our two experiments. **(B)** Fixed trolley in the anterior-posterior direction represents the set-up used by previous studies. The anterior-posterior displacements of trolley (blue line), frame (red line) as well as their anterior-posterior displacement differences (yellow line) in free **(C)** and fixed **(D)** trolley conditions. The magnitudes of anterior-posterior spring forces during normal walking in free **(E)** and fixed **(E)** trolley conditions. The magnitudes of anterior-posterior spring forces during walking with extra movement in the anterior-posterior direction in free **(G)** and fixed **(H)** trolley conditions.

### The estimation of frictions between waist belt and sliders as well as between cart and trolley

To estimate the friction for transverse rotation in the "Free" condition, and the friction of movement in the AP direction, we embedded an artificial pelvis (weight = 2.0 kg) inside the waist belt of the frame which was used in experiment 2. We placed a force transducer at the top of artificial pelvis. The transducer provided 3D forces and moments. Three kinematic markers were also placed on the back side of force transducer. Using a vertical rod connected to the transducer, the researcher slowly pushed and pulled the transducer toward the forward and backward directions together with imposing rotations. These imposed movements caused an anterior-posterior pelvis displacement and a transverse pelvis rotation, respectively (**Fig S2. A**).

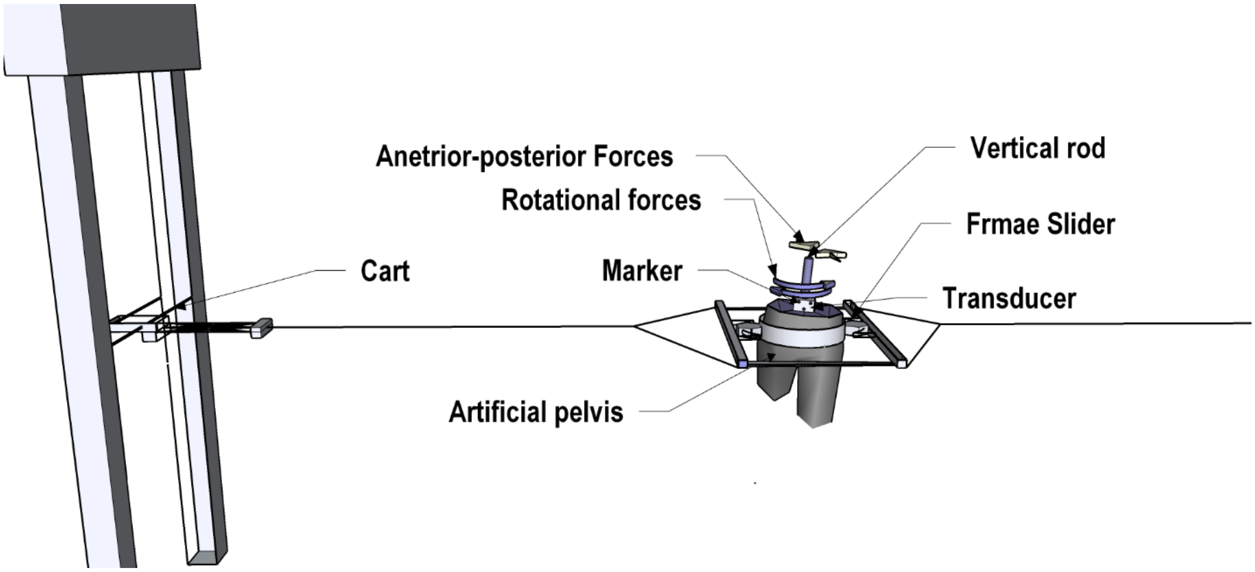


**Fig S2.** Schematic representation of experimental set up and imposed forces induced by researcher on artificial pelvis.

Like in the above described measurements, the results showed that the anterior-posterior force was <8N, indicating that the friction between trolley and cart is low **(Fig S3. B).** Interestingly, these forces appeared to be mostly displacement **(Fig S3. B)** and acceleration **(Fig S3. D)** dependent, suggesting that actual friction played a minor role.

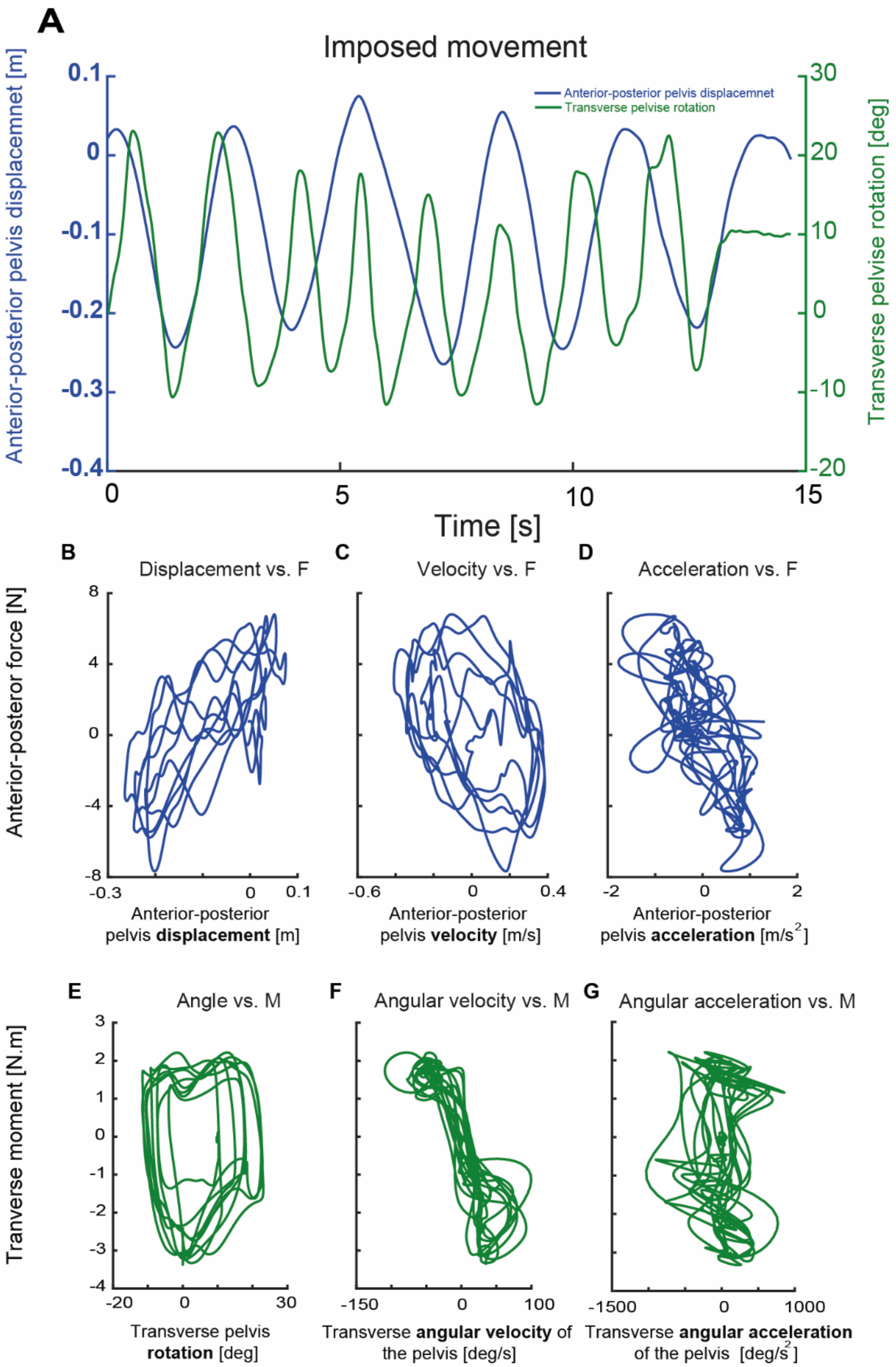


**Fig S3. (A)** the linear and angular changes of the artificial pelvis (blue line represents anterior-posterior pelvis displacement and green line represents transverse pelvis rotation). The relationship between anterior-posterior force and anterior-posterior pelvis displacement **(B)**, velocity **(C)**, and acceleration **(D)** during the imposed anterior-posterior forces on the artificial pelvis. The relationship between transverse moment and transverse pelvis rotation **(E)** as well as transverse angular velocity **(F)** and **(G)** acceleration of the artificial pelvis during the imposed rotational forces.

Our analyses on imposed rotational forces showed that a transverse moment of around 2 Nm was sufficient to overcome the friction between the waist belt and the frame sliders (**Fig S3. E-G**).

It can be concluded that the frictions between the trolley and cart as well as between the waist belt and frame sliders are minimal.
